# Decomposing complex links between the childhood environment and brain structure in school-aged youth

**DOI:** 10.1101/2020.04.28.063461

**Authors:** Seok-Jun Hong, Lucinda Sisk, Camila Caballero, Anthony Mekhanik, Amy K. Roy, Michael P. Milham, Dylan G. Gee

**Affiliations:** Center for the Developing Brain, Child Mind Institute, New York, NY, USA; Center for Neuroscience Imaging Research, Institute for Basic Science, Sungkyunkwan University, Suwon, South Korea; Department of Biomedical Engineering, Sungkyunkwan University, Suwon, South Korea; Department of Psychology, Yale University, New Haven, CT, USA; Department of Psychology, Fordham University, Bronx, NY, USA; Center for Biomedical Imaging and Neuromodulation, Nathan Kline Institute, Orangeburg, NY, USA

**Keywords:** brain development, adversity, environment, childhood, neuroanatomy, subtyping

## Abstract

Childhood experiences play a profound role in conferring risk and resilience for brain and behavioral development. However, how different facets of the environment shape neurodevelopment remains largely unknown. Here we sought to decompose heterogeneous relationships between environmental factors and brain structure in 989 school-aged children from the Adolescent Brain Cognitive Development Study. We applied a cross-modal integration and clustering approach called ‘Similarity Network Fusion’, which combined two brain morphometrics (*i*.*e*., cortical thickness and myelin-surrogate markers), and key environmental factors (*i*.*e*., trauma exposure, neighborhood safety, school environment, and family environment) to identify homogeneous subtypes. Depending on the subtyping resolution, results identified two or five subgroups, each characterized by distinct brain structure-environment profiles. Notably, more supportive caregiving and school environments were associated with increased myelination, whereas less supportive caregiving, higher family conflict and psychopathology, and higher perceived neighborhood safety were observed with increased cortical thickness. These subtypes were highly reproducible and predicted externalizing symptoms and overall mental health problems. Our findings support the theory that distinct environmental exposures differentially influence neurodevelopment. Delineating more precise associations between risk factors, protective factors, and brain development may inform approaches to enhance risk identification and optimize interventions targeting specific experiences.

## INTRODUCTION

Experiences during childhood play a crucial role in shaping the developing brain, behavior, and risk for psychopathology (Chen & Baram, 2016; Gee, 2016; McLaughlin et al., 2017; Nelson & Gabard-Durnam, 2020; Opendak et al., 2017; Tottenham, 2012). A nuanced understanding of how early experiences alter structural brain development is critical to elucidating the mechanisms by which childhood adversity confers risk for psychopathology, and protective environmental factors buffer that risk. Early adverse experiences have been shown to disrupt neurodevelopment on a cellular level (Abbink et al., 2019; Bath et al., 2016; Bordner et al., 2011; Johnson & Kaffman, 2018), and a growing literature has identified alterations in structural brain features such as gray matter volume (De Bellis et al., 1999; Hair et al., 2015; Hanson et al., 2012; Hodel et al., 2015; Kribakaran et al., 2020; Mackes et al., 2020; McEwen, 2016; Noble et al., 2015; Sheridan et al., 2012; Teicher et al., 2016; Tottenham et al., 2010), cortical thickness (Gold et al., 2016; Kelly et al., 2013; Lim et al., 2018; McLaughlin et al., 2014; Monninger et al., 2019), white matter tract integrity (Bick et al., 2015; Hanson et al., 2013; Ho et al., 2017; Howell et al., 2013; Kircanski et al., 2019), and myelination (Bath et al., 2016; Bordner et al., 2011; Juraska & Kopcik, 1988; Makinodan et al., 2012) following adversity.

Much of the existing knowledge about environmental influences on brain development has stemmed from research focusing on a single type of experience (*e*.*g*., physical abuse, neglect, exposure to violence) or aggregating across different types of exposures to adversity (De Bellis et al., 1999; Mehta et al., 2009; Tomoda et al., 2009, 2012). While such evidence has been foundational in establishing the deleterious effects of early adversity, there is vast heterogeneity in both the nature of adversity exposure and in outcomes (Cohodes et al., 2020). The frequent co-occurrence of adverse experiences (Green et al., 2010) and additional complexity of family, neighborhood, and school environments present further challenges to precisely linking environmental factors with variation in brain structure.

Dimensional approaches have increasingly focused on key aspects of early adversity (Cicchetti & Toth, 1995; Cohodes et al., 2020; Everaerd et al., 2016; McCoy, 2013; McLaughlin et al., 2014; Pynoos et al., 1999), including the type of adversity experienced (Dennison et al., 2019; Machlin et al., 2019; McLaughlin et al., 2014; Miller et al., 2018; Sheridan et al., 2017). Previous work directly comparing distinct types of exposures (e.g., physical abuse, sexual abuse, physical neglect, emotional neglect) has demonstrated differential impacts on brain structure (Cassiers et al., 2018; Edmiston et al., 2011; Heim et al., 2013; Tomoda et al., 2009, 2012; van Harmelen et al., 2010). Examining findings across studies of specific types of adversity has also suggested unique associations with brain structure. For example, distinct regional patterns of reduced cortical thickness have been observed among children exposed to severe neglect in institutional care (Hodel et al., 2015; McLaughlin et al., 2014) versus children exposed to abuse (Busso et al., 2017; Gold et al., 2016; Lim et al., 2018).

While much of the literature on environmental influences has focused on adversity, a growing body of research has examined normative variation in environmental factors. The relationship between child and primary caregivers is thought to be particularly influential in shaping neurodevelopment (Tottenham, 2018; Gee et al., 2016), with longitudinal evidence that positive and negative parenting behaviors are associated with differential change in brain development in adolescents (Whittle et al., 2014, 2016). Positive, more sensitive parenting has been found to predict accelerated cortical thinning in the anterior cingulate in males, and orbitofrontal cortex (Whittle et al., 2014), as well as increased volume in the posterior insular cortex (Matsudaira et al., 2016) and across the whole brain (Kok et al., 2015, 2018). Negative, more aggressive parenting has been shown to predict increased thickening of the superior frontal gyrus and lateral parietal lobe in males (Whittle et al., 2016), and has been associated with larger anterior cingulate and orbitofrontal cortex volumes (Whittle et al., 2009). While null effects of caregiving on cortical thickness have also been reported (Avants et al., 2015; Leblanc et al., 2017), accumulating evidence highlights the importance of the caregiver/child relationship and demonstrates that both positive and negative caregiving experiences impact structural brain development (Deane et al., 2020). Factors such as greater neighborhood disadvantage (Whittle et al., 2017) and positive school environments (Piccolo et al., 2019) have also been independently associated with increases in cortical thickness during development. However, less is known about the ways in which neighborhood and school contexts may interact with other environmental factors to influence brain structure.

Variations in cortical thickness and volume have been widely studied in the literature and may reflect processes such as synaptic pruning and remodeling (Huttenlocher, 1979; Huttenlocher et al., 1982; Huttenlocher & Dabholkar, 1997) or stress-induced neuronal atrophy (Horchar & Wohleb, 2019; Wellman et al., 2020). Though less studied in humans, myelination is thought to increase throughout development (Lebel & Deoni, 2018) and is sensitive to adversity in rodent models (Bordner et al., 2011; Carlyle et al., 2012; Makinodan et al., 2017). However, no studies to our knowledge have investigated differential effects of early environmental exposure type on cortical thickness and myelination. As both of these processes undergo marked maturational changes during childhood (Dean et al., 2015; Lyall et al., 2015) and have been implicated in various psychopathologies that often emerge during development (Hanford et al., 2016; Norbom et al., 2019; Schmaal et al., 2017; van Erp et al., 2018), understanding how cortical thickness and myelination are shaped by specific aspects of early environments is an important gap to address.

Given the complexity of associations between early experiences and brain development, multivariate approaches capable of handling high-dimensional data show promise for elucidating associations between adversity and brain structure. One such data-driven approach is subtyping, which aims to identify subgroups of individuals with similar neural and behavioral characteristics, and to examine differential outcomes between these subgroups. This approach has been applied effectively in studies examining subtypes of individuals with psychiatric disorders (Fair et al., 2012; Hong et al., 2018, 2019; Sun et al., 2015), but not yet within the context of the childhood environment and brain development.

In the current study, we aimed to decompose heterogeneous relationships between specific environmental exposures and brain structure during development. To address this goal, we leveraged a novel multimodal data integration framework, similarity network fusion (SNF) (Wang et al., 2014), and applied it to openly shared, large-scale data derived from the Adolescent Brain Cognitive Development (ABCD) Study (Casey et al., 2018). Compared to traditional unimodal approaches, this framework allowed us to take into account environmental factors and structural brain features simultaneously in clustering, thus unveiling differential subtypes that are more readily interpretable in both environmental and neurobiological domains. Following SNF, we then tested the validity of those subtypes by predicting clinical symptoms (which were not used for subtyping) based on brain imaging features of each subtype in a replication dataset. We hypothesized that differential patterns of myelin and cortical thickness would be associated with discrete measures indexing the childhood environment, resulting in subtypes representing co-occurrence of specific structural variation and environmental exposures. By exploring more precise associations between environmental exposures and structural variation in a large, population-based, demographically diverse sample, we aim to enhance understanding of how environmental and brain structural variation co-occur and relate to mental health during childhood.

## METHODS AND MATERIALS

### Subjects

Participants in our study were 989 school-aged youth (9-10 years old), whose data were obtained from the Adolescent Brain Cognitive Development (ABCD) Study (Casey et al., 2018). This ongoing project aims to recruit over 11,000 children from 21 different sites based on harmonized protocols (Casey et al., 2018) and follow them over ten years to comprehensively characterize psychological and neurobiological development from pre-adolescence to young adulthood. Parents provided written informed consent, and children provided verbal assent for study participation. Full details of ethics and oversight in the ABCD Study have been previously published (Clark et al., 2018). We applied a set of inclusion and exclusion criteria to select a subsample from this broader pool of participants: *i*) included only participants with all data of interest, including T1- and T2-weighted MRI scans indexing brain structure, 8 phenotypic scores related to environmental conditions for youth, and 3 scores indexing different facets of mental health (see Environmental data and Clinical data), *ii*) if there were siblings, only the oldest child in each family was included, and *iii*) excluded those participants with diagnoses of autism spectrum disorder or epilepsy. Apart from these criteria, participants affected by the error related to structural MRI data reported in Known Issues with Data Release 2.0 (https://nda.nih.gov/edit_collection.html?id=2573) were excluded. Finally, we excluded participants with a lower quality of MRI and preprocessing results based on the ABCD Study’s FreeSurfer quality control conducted by trained technicians to identify data showing evidence of excessive motion, pial overestimation, white matter underestimation, inhomogeneity, or artifacts (Hagler et al., 2019). Based on these criteria and external quality control screening, there were 2379 remaining participants, collected across 13 sites and 2 different scanners. To reduce the computational cost in processing such high-volume data, we randomly selected 1000 participants for inclusion in the present study. We internally performed quality control procedures on these 1000 participants’ data, which consisted of visual inspection of remaining cases for the FreeSurfer processing derivatives (*e*.*g*., cortical surface) as well as the z-scores of imaging features to identify outliers (see later paragraphs for details of quality control processes). This internal quality control procedure excluded 11 participants, resulting in a final n=989. The final sample was randomly split into 495 discovery and 494 replication cases. Among these, 230/213 participants (discovery/replication) were drawn from the first release of the ABCD Study (NIMH Data Archive Release 1.1, DOI: 10.15154/1412097) and 265/281 participants (discovery/replication) from the second release of the ABCD Study (DOI: 10.15154/1503209). All details of this participant sampling process are summarized in the **Supplementary Material and Table 1**. The discovery and replication samples did not show differences in any demographic data including age, sex, data collection site, race, ethnicity, parental education, and household income. These profiles are reported in **Table 1**. Further details about the brain imaging and behavioral data collected in the ABCD Study can be found in the original data descriptor papers (Barch et al., 2018; Casey et al., 2018).

**Table 1.**
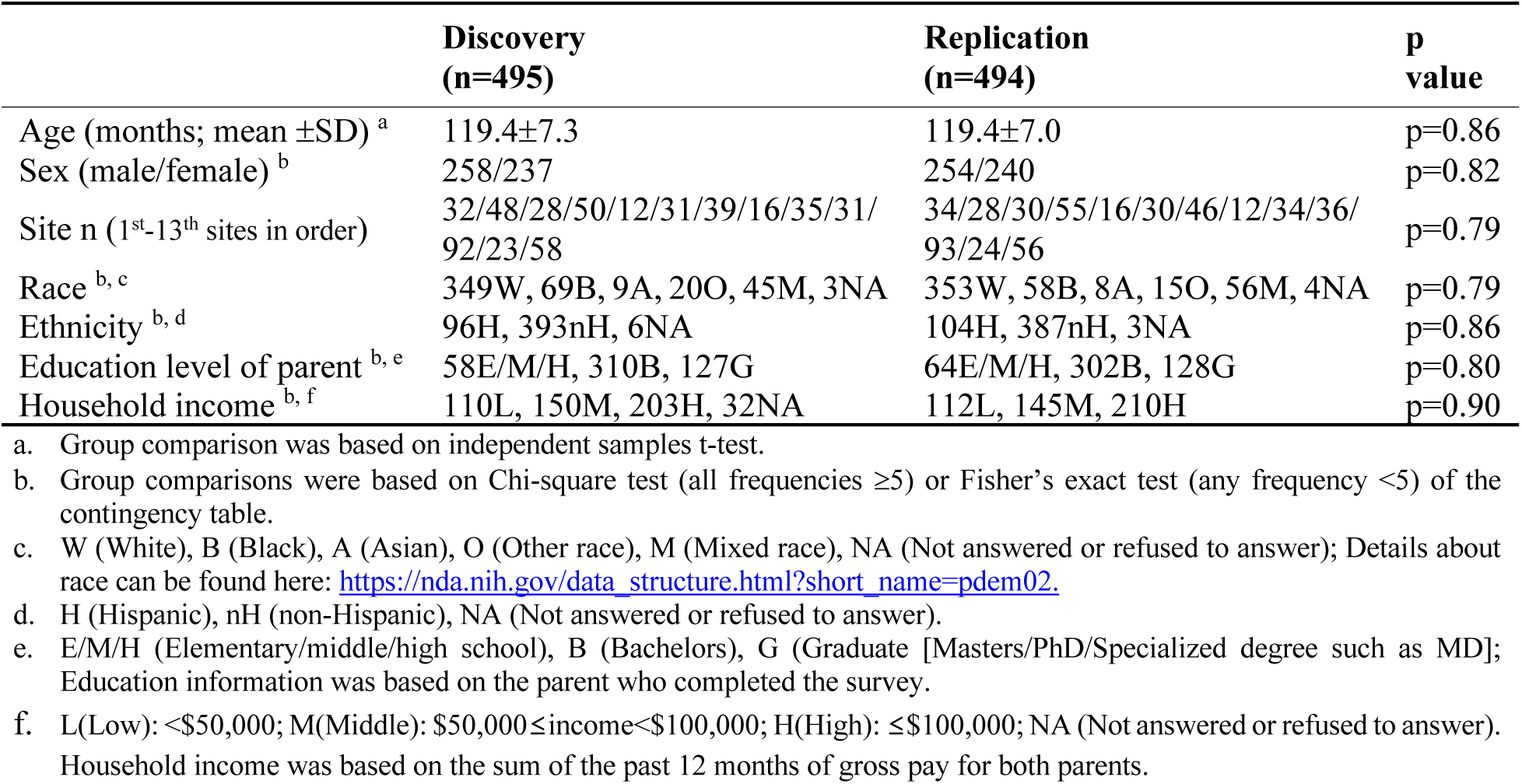
Demographic characteristics of included participants

|  | **Discovery**<br>**(n=495)** | **Replication**<br>**(n=494)** | **p**<br>**value** |
| --- | --- | --- | --- |
| Age (months; mean ±SD) <sup>a</sup> | 119.4±7.3 | 119.4±7.0 | p=0.86 |
| Sex (male/female) <sup>b</sup> | 258/237 | 254/240 | p=0.82 |
| Site n (1<sup>st</sup>-13<sup>th</sup> sites in order) | 32/48/28/50/12/31/39/16/35/31/<br>92/23/58 | 34/28/30/55/16/30/46/12/34/36/<br>93/24/56 | p=0.79 |
| Race <sup>b, c</sup> | 349W, 69B, 9A, 20O, 45M, 3NA | 353W, 58B, 8A, 15O, 56M, 4NA | p=0.79 |
| Ethnicity <sup>b, d</sup> | 96H, 393nH, 6NA | 104H, 387nH, 3NA | p=0.86 |
| Education level of parent <sup>b, e</sup> | 58E/M/H, 310B, 127G | 64E/M/H, 302B, 128G | p=0.80 |
| Household income <sup>b, f</sup> | 110L, 150M, 203H, 32NA | 112L, 145M, 210H | p=0.90 |

a. Group comparison was based on independent samples t-test.
b. Group comparisons were based on Chi-square test (all frequencies $\geq 5$ ) or Fisher's exact test (any frequency $< 5$ ) of the contingency table.
c. W (White), B (Black), A (Asian), O (Other race), M (Mixed race), NA (Not answered or refused to answer); Details about race can be found here: https://nda.nih.gov/data_structure.html?short_name=pedem02.
d. H (Hispanic), nH (non-Hispanic), NA (Not answered or refused to answer).
e. E/M/H (Elementary/middle/high school), B (Bachelors), G (Graduate [Masters/PhD/Specialized degree such as MD]); Education information was based on the parent who completed the survey.
f. L(Low): $< \$50,000$ ; M(Middle): $\$50,000 \leq \text{income} < \$100,000$ ; H(High): $\leq \$100,000$ ; NA (Not answered or refused to answer). Household income was based on the sum of the past 12 months of gross pay for both parents.

### Imaging data

Structural imaging data consisted of T1-weighted (T1w) and T2-weighted (T2w) MRI, both acquired using a 3T Siemens Prisma scanner. Specifically, the T1w acquisition was based on a 3D inversion prepared RF-spoiled gradient echo scan (TE=2.88ms, TR=2500ms, flip angle = 8°, 1mm isotropic voxels, 2x parallel imaging) using prospective motion correction (Tisdall et al., 2012; White et al., 2010). The T2w acquisition was carried out based on a 3D T2w variable flip angle fast spin echo sequence (TE=565ms, TR=3200ms, 1mm isotropic voxels, 2x parallel imaging), also with prospective motion correction.

### Environmental data

Environmental factors were selected to characterize youth environment with regard to trauma exposure, caregiver behaviors, family functioning, neighborhood safety, and school environment. Six of these factors were assessed using measures from the ABCD Culture and Environment Sum Scores: neighborhood safety (mean score of the ABCD Parent Neighborhood Safety/Crime Survey Modified from PhenX), school environment (‘school environment’ subscale from the ABCD School Risk and Protective Factors Survey), parental support (mean score of first five items from the ABCD Children’s Report of Parental Behavioral Inventory), caregiver support (mean score of second five items from the ABCD Children’s Report of Parental Behavioral Inventory), parental monitoring (mean score of the ABCD Parental Monitoring Survey), and family conflict (‘family conflict’ subscale from the ABCD Youth Family Environment Scale-Family Conflict Subscale Modified from PhenX) (Hoffman et al., 2019). Trauma exposure was assessed by computing a summed score across the 17 categories queried in the Traumatic Events measure of the ABCD Parent Diagnostic Interview for DSM-5 [Kiddie Schedule for Affective Disorders and Schizophrenia (KSADS) (Kaufman et al., 1997)]. This section of the semi-structured interview assesses events such as physical and sexual abuse, exposure to domestic violence, and exposure to community violence. Family history of mental health problems was assessed via the sum of endorsements for substance use, criminal activities, and mental health concerns across all immediate family members (mother, father, full siblings) queried in the ABCD Family History Assessment. Further details on all eight environmental measures can be found in the **Supplementary Material**.

### Clinical data

Three measures of psychiatric symptoms (T-scores indexing symptoms of internalizing problems, externalizing problems, and total problems) were selected from the parent-reported Child Behavior Checklist (CBCL; Achenbach & Rescorla, 2001) to evaluate clinical profiles of the identified subtypes.

### Image preprocessing and feature extraction

The acquired individual T1w and T2w MRI data underwent an established processing pipeline from the Human Connectome Project (HCP) (Glasser et al., 2013). This contains multiple optimized preprocessing steps for cortical surface extraction and volume/surface alignment processes, details of which can be found in the original pipeline paper (Glasser et al., 2013). Briefly, this pipeline consists of three main stages: *i)* the ‘PreFreeSurfer’ stage to produce an undistorted native structural volume space for each participant, align the T1w and T2w images, perform a bias-field correction, and register the participant’s native structural volume space to MNI space, *ii)* the ‘FreeSurfer’ stage (using the version 5.3 FreeSurfer software) to segment the volume into predefined structures, reconstruct white and pial cortical surfaces, and perform FreeSurfer’s standard folding-based surface registration to their surface atlas, and finally *iii)* the ‘PostFreeSurfer’ stage to produces all necessary NIFTI volume and GIFTI surface files, along with applying the surface registration and creating the final brain mask.

To probe heterogeneous relationships between environmental conditions and brain development, we analyzed two widely employed structural MRI features, namely cortical thickness and T1w/T2w ratio indexing myelination. Cortical thickness has been associated with multiple cellular features that are closely related to neurodevelopment including neuropil volume (Schüz & Palm, 1989), neuronal density (Collins et al., 2010; la Fougère et al., 2011), arborization (Scholtens et al., 2014), and intracortical connectivity (Wagstyl & Lerch, 2018). Myelination is an additional biological process that occurs throughout development, and supports neuronal adaptation during co-occurring processes such as synaptogenesis and pruning (Silbereis et al., 2016). In our study, myelination was indexed by a measure automatically extracted in the second ‘FreeSurfer’ stage of the above HCP pipeline, measured as the distance of corresponding vertices between the white and pial boundary. This myelin-surrogate marker was constructed in the third ‘Post-FreeSurfer’ stage by dividing T1w intensity by T2w intensity (thus, a T1w/T2w ratio) at each cortical point (vertex) across the whole brain (Glasser & Van Essen, 2011). Notably, while the original cortical surfaces that the HCP pipeline generated had 32,000 vertices per hemisphere, we opted to downsample this surface mesh to 10,242 vertices at each hemisphere in order to reduce the computational cost in the following subtype analyses.

Once the feature preprocessing was complete, we visually inspected the reconstructed surfaces and cortical masks across individual brains for quality control. We also performed quantitative outlier detection based on vertex-wise z-score maps of both cortical thickness and myelin. Participants were excluded if >20% of the vertices in both features had z-scores greater than 3.09 (a threshold corresponding to the p-value 0.001). This provided a final sample for discovery (n=495) and replication (n=494) data in the current study.

### Feature preprocessing

The above feature extraction resulted in two sets of 20,484 feature values (from the cortical thickness and myelin maps) as well as 8 phenotypic and 3 clinical scores for each individual. To control for possible confounds (*i*.*e*., age, sex, data collection site), we performed *i)* a statistical correction for age and sex effects on imaging features (Hong et al., 2018) and *ii)* ComBat harmonization (*i*.*e*., Combining Batch effects, a Bayesian approach based on a linear model to estimate and remove site batch effects)(Fortin et al., 2017), for both imaging and phenotypic scores. We then normalized each feature using z-scoring.

### Subtyping based on similarity network fusion (Figure 1)

Subtyping was performed using similarity network fusion (SNF), an algorithm originally developed to integrate multimodal data in the genetic field (*e*.*g*., microRNA expression and DNA methylation) (Wang et al., 2014). This approach has recently been applied to neuroimaging studies to objectively subtype brain structures across transdiagnostic samples (Stefanik et al., 2018). SNF was performed on the measures of brain structure and environmental factors; the measures of clinical symptoms were not included in any step of SNF. Subtyping using SNF consisted of the following four steps:

**Figure 1.**
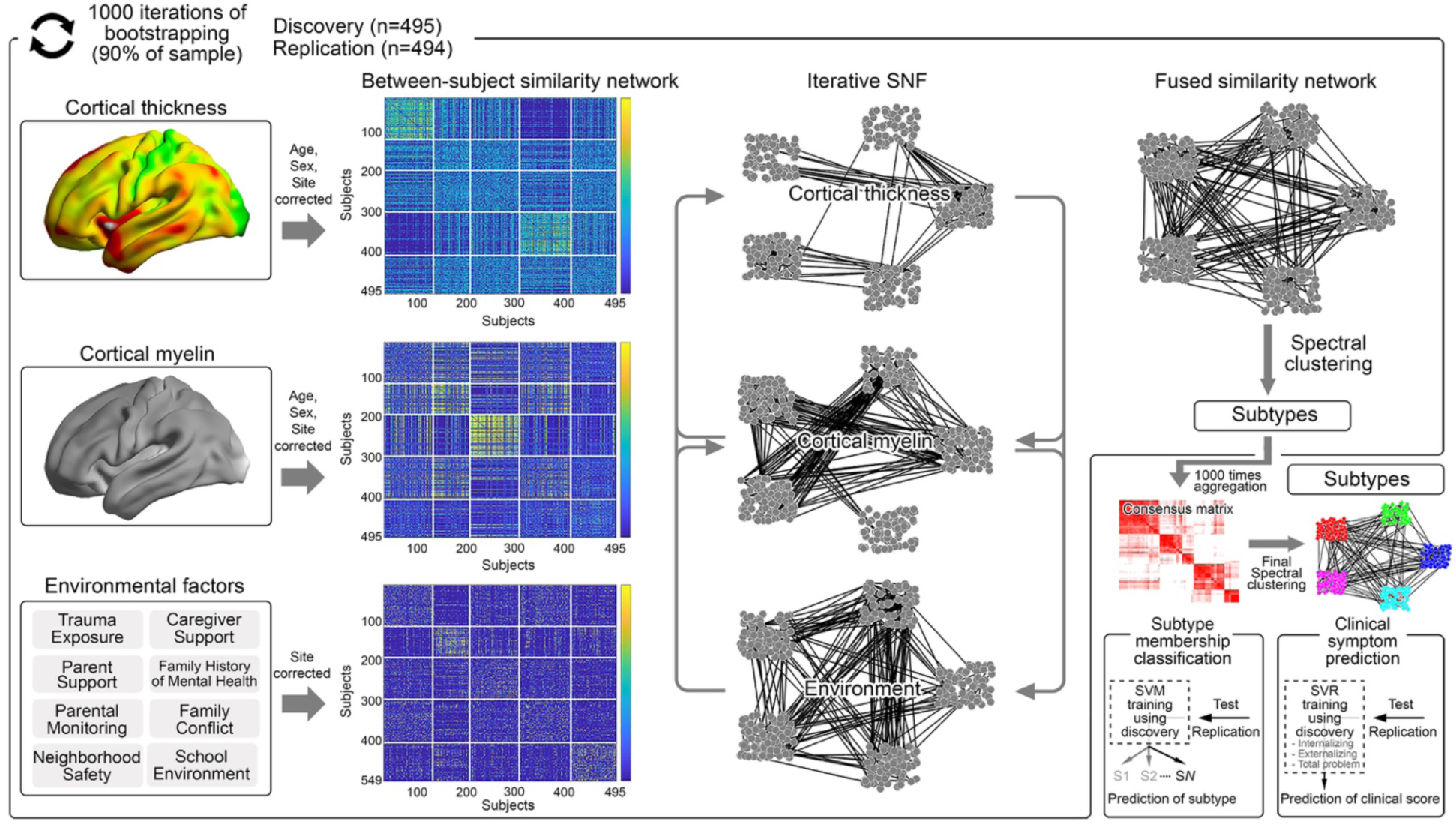
General method for subtyping based on Similarity Network Fusion (SNF). Two brain structural features and eight environmental factors were used to create a between-subjects similarity network at each modality. The resulting similarity graphs were then entered into SNF which performs iterative non-linear fusion processes to combine them, resulting in a single ‘fused similarity network’. Spectral clustering was then applied to find homogeneous subgroups. To find the most reproducible clustering results, we bootstrapped 90% of the samples and repeated the above SNF subtyping. This procedure was iterated 1000 times, and the subtype results were aggregated to construct a consensus clustering matrix. The final clustering result was obtained based on spectral clustering of the consensus matrix. The optimal clustering solution was determined by an established consensus index (Monti et al., 2003). To assess the significance of identified subtypes, two prediction analyses were conducted of subtype membership classification and clinical symptoms. These analyses aimed to demonstrate both generalizability and utility in predicting clinical symptoms using independent samples.

#### i) Calculation of unimodal affinity matrices

We first computed three between-subjects distance matrices for cortical thickness, T1w/T2w ratio maps, and the environmental phenotypic scores separately. The distance matrix was calculated consistently across the features based on the ‘cosine similarity’ kernel, which was then used to generate between-subjects affinity matrices (each cell representing the similarity of distance profiles between two given participants) using a scaled exponential similarity kernel. The mathematical details of this kernel can be found in the original algorithm paper (Wang et al., 2014).

#### ii) Fusion of multimodal affinity matrices

The resulting three between-subjects affinity matrices (each from cortical thickness, myelin, and environmental phenotypic scores) were then input to SNF to generate a single fused affinity matrix. Three parameters were employed in SNF (*i*.*e*., *k*: number of nearest neighbors used to fuse the affinity matrices [how many local neighbors used to calculate the between-subjects similarity in SNF], T: number of iterations in SNF algorithms, and μ: hyperparameters related to the scaling process of each feature). We selected those parameters as the original paper recommended (k=30, T=20, μ=0.5) (Wang et al., 2014). As detailed further below, these parameters resulted in the most reproducible subtype findings between the discovery and replication datasets.

#### iii) Spectral clustering to identify subtypes

The generated single fused affinity matrix was input to the spectral clustering algorithm to identify homogeneous subtypes. Of note, to obtain more reproducible and outlier-robust subtype findings, we performed the above SNF process 1000 times iteratively, based on the bootstrapped samples (90% of cases resampled without replacement) and constructed a consensus matrix, varying the clustering number (*C*) from 2 to 20. Each individual cell of this consensus matrix represents how consistently a given pair of participants was grouped together among 1000 iterations at a given clustering number (2-20).

#### iv) Clustering solution evaluation

We used a previously established approach, ‘cumulative consensus distribution’, to determine the clustering solution. Detailed information regarding the mathematical principle and motivation can be found in the original paper (Monti et al., 2003). Briefly, systematically evaluating *C* from 2 to 20, this approach evaluated up to which value *C* increased the degree of consensus for the clustering solution, compared to the previous *C* (*i*.*e*., *C*- 1). Using this criterion, we selected the *C* with the highest subtyping stability across differently sampled cases.

Importantly, the entire subtyping process (*i-iv*) was performed two separate times for the discovery and replication datasets to independently assess the reproducibility of findings.

### Subtype profiling

The identified subtypes were then evaluated in terms of brain and environmental features. For quantitative evaluation, we performed analysis of covariance (ANCOVA) on cortical thickness, myelin maps, and environmental phenotypic scores for main group effects (*i*.*e*., subtypes) while statistically correcting for age, sex, and data collection site. The family-wise error due to multiple comparisons for phenotypic scores was controlled by the false discovery rate (FDR; Benjamini & Hochberg, 1995). For brain features, we additionally included the global mean (*i*.*e*., whole brain averaged cortical thickness and T1w/T2w ratio) as a nuisance variable to highlight region-specific subtype differences across the brain. To focus on only reproducible findings, we mapped significant clusters from ANCOVA that overlapped between the discovery and replication datasets. For these overlapping regions, we evaluated their spatial patterns and profiles of cortical thickness and T1w/T2w ratio across subtypes. The family-wise error due to massive univariate vertex-wise multiple comparisons was controlled by random field theory (RFT; Worsley et al., 1999) at 0.05 (cluster-defining threshold=0.025). Beyond the regional effects, we also performed ANCOVA on the whole-brain mean values, as they reflect global effects from more diffuse biological substrates.

### Prediction analysis

To validate our subtype results, we carried out two prediction analyses: *i)* subtype classification and *ii)* prediction of clinical symptoms for each subtype. To ensure the generalizability of our findings, the training of each analysis was conducted based on the discovery dataset, whereas the test was based on the independent replication dataset.

#### i) Classification of subtype membership

This analysis aimed to assess how generalizable the brain and environmental phenotypic profiles of identified subtypes were to unseen cases. To do this, we trained a multiclass support vector machine algorithm by using the full imaging features (2×20,484 for cortical thickness and myelin maps) and 8 environmental phenotypic scores of the 495 discovery cases as predictors and their subtype information as a responder. The accuracy of the classification was measured by entering the same imaging and phenotypic features of the 494 unseen replication cases into the trained classifier and comparing the predicted subtype with that which resulted from the SNF analysis based on the entire 989 cases.

#### ii) Prediction of clinical symptoms

Our second prediction analysis sought to associate the brain imaging features with clinical symptoms (*i*.*e*., internalizing, externalizing, and total problem) for each identified subtype. Of note, these clinical variables were not used in the SNF subtyping. We trained the support vector regression for each subtype based on brain imaging features of the discovery cases to predict their clinical symptom scores. For feature selection, we performed a linear regression between each feature and targeted clinical symptom scores of the discovery dataset and entered only those showing a significant correlation (|t|>2) into the classifier to reduce the feature dimensionality and optimize the training quality. We then tested this subtype-specific classifier using the independent replication dataset, entering the same brain imaging features. The prediction accuracy was measured by a non-parametric Spearman correlation between the original clinical symptom scores and the predicted scores across internalizing, externalizing symptoms, and total problems. To test the utility of our subtyping approach, we also conducted the same prediction of clinical symptom scores but without subtype information. In other words, we trained the classifier using the 495 discovery cases and tested it based on the 494 replication cases. This analysis aimed to address whether identifying subgroups provides a more homogeneous predictor-responder relationship, thereby improving prediction accuracy.

### Data and code availability

All data analyzed in this study are accessible through an official request to the NIMH Data Archive (https://nda.nih.gov/abcd). All other relevant materials (*e*.*g*., a participant list) and code used for data sampling, SNF, and statistical analyses are publicly available at https://github.com/Yale-CANDLab/ABCD_Env_Subtyping.

## RESULTS

### Subtyping based on SNF

Based on the area under the curve for a cumulative distribution function (CDF) of consensus values (**Supplementary Figure 1B**), we focused on the subtyping results at clustering numbers *C*=2 and *C*=5, given that these *C*s made the largest increases (ΔCDF>0) of consensus degree in both the discovery and replication datasets. Indeed, for these solutions, high stability of between-subject subgroups across bootstrapped samples was observed in the consensus matrices (**Supplementary Figure 1A**), and the subtyping patterns were highly reproducible in the replication data (**Figure 2 and 3**). Notably, resulting subtypes in the discovery dataset did not show any differences in age, sex, site, race, ethnicity, parental education, or annual household income in both the *C*=2 and *C*=5 solutions (**Table 2;** see also **Supplementary Table 2** for the replication dataset in which there were no differences for any variables except for household income).

**Table 2.** Demographic profiles of 2- and 5-subtype solutions (Discovery)

| 2-subtype profiling | Subtype 1<br>(n=258) | Subtype 2<br>(n=237) | p value |
| --- | --- | --- | --- |
| Age (months; mean ±SD) <sup>a</sup> | 119.7±7.3 | 119±7.3 | p=0.27 |
| Sex (male/female) <sup>b</sup> | 136/122 | 122/115 | p=0.75 |
| Site n (1<sup>st</sup>-13<sup>th</sup> sites in order) <sup>b</sup> | 14/23/13/23/5/15/19/10/15/19/<br>53/11/38 | 18/25/15/27/7/16/20/6/20/12/<br>39/12/20 | p=0.60 |
| Race <sup>b, c</sup> | 190W, 36B, 4A, 8O, 20M | 159W, 33B, 5A, 12O, 25M, 3NA | p=0.26 |
| Ethnicity <sup>b, d</sup> | 55H, 200nH, 3NA | 41H, 19nH, 3NA | p=0.26 |
| Education level of parent <sup>b, e</sup> | 31E/M/H, 168B, 59G | 27E/M/H, 142B, 68G | p=0.33 |
| Household income <sup>b, f</sup> | 58L, 80M, 104H, 16NA | 52L, 70M, 99H, 16NA | p=0.97 |

| 5-subtype profiling | Subtype 1<br>(n=82) | Subtype 2<br>(n=106) | Subtype 3<br>(n=112) | Subtype 4<br>(n=93) | Subtype 5<br>(n=102) | p value |
| --- | --- | --- | --- | --- | --- | --- |
| Age (months; mean ±SD) <sup>a</sup> | 119.4±7.4 | 119.9±7.5 | 119.8±7.6 | 118.8±6.9 | 118.9±7.1 | p=0.71 |
| Sex (male/female) <sup>b</sup> | 39/43 | 51/55 | 64/48 | 51/42 | 53/49 | p=0.60 |
| Site n (1<sup>st</sup>-13<sup>th</sup> sites in order) <sup>b</sup> | 8/8/6/7/2/4/<br>5/2/7/5/13/<br>4/11 | 4/11/8/10/2/<br>8/9/1/8/5/19/<br>4/17 | 11/9/5/13/1/<br>8/12/3/9/8/1<br>8/4/11 | 2/7/4/13/3/4/<br>8/6/3/10/24/<br>2/7 | 7/13/5/7/4/7/<br>5/4/8/3/<br>18/9/12 | p=0.75 |
| Race <sup>b, c</sup> | 60W, 12B,<br>3A, 6M,<br>1NA | 81W, 15B,<br>1A, 5O, 4M | 72W, 15B,<br>2A, 7O,<br>15M, 1NA | 68W, 13B,<br>2A, 1O, 9M | 68W, 14B,<br>1A, 7O,<br>11M, 1NA | p=0.39 |
| Ethnicity <sup>b, d</sup> | 15H, 66nH,<br>1NA | 25H, 79nH,<br>2NA | 21H, 91nH | 18H, 74nH,<br>1NA | 17H, 83nH,<br>2NA | p=0.77 |
| Education level of parent <sup>b, e</sup> | 6E/M/H,<br>57B, 19G | 16E/M/H,<br>64B, 26G | 14E/M/H,<br>69B, 29G | 11EMH/61B<br>/21G | 11EMH/59B<br>/32G | p=0.69 |
| Household income <sup>b, f</sup> | 22L, 24M,<br>29H, 7NA | 25L, 32M,<br>43H, 6NA | 21L, 34M,<br>47H, 10NA | 18L, 32M,<br>37H, 6NA | 24L, 28M,<br>47H, 3NA | p=0.81 |

a. Group comparison was based on independent samples t-test.
b. Group comparisons were based on Chi-square test (if all frequencies $\geq 5$ ) or Fisher's exact test (if any frequency $< 5$ ) of the contingency table.
c. W (White), B (Black), A (Asian), O (Other race), M (Mixed race), NA (Not answered or refused to answer); Details about race can be found here: https://nda.nih.gov/data_structure.html?short_name=pdem02.
d. H (Hispanic), nH (non-Hispanic), NA (Not answered or refused to answer).
e. E/M/H (Elementary/middle/high school), B (Bachelors), G (Graduate [Masters/PhD/Specialized degree such as MD]); Education information was based on the parent who completed the survey.
f. L(Low): $< \$50,000$ ; M(Middle): $\$50,000 \leq \text{income} < \$100,000$ ; H(High): $\leq \$100,000$ ; NA (Not answered or refused to answer). Household income was based on the sum of the past 12 months of gross pay for both parents.

**Figure 2.**
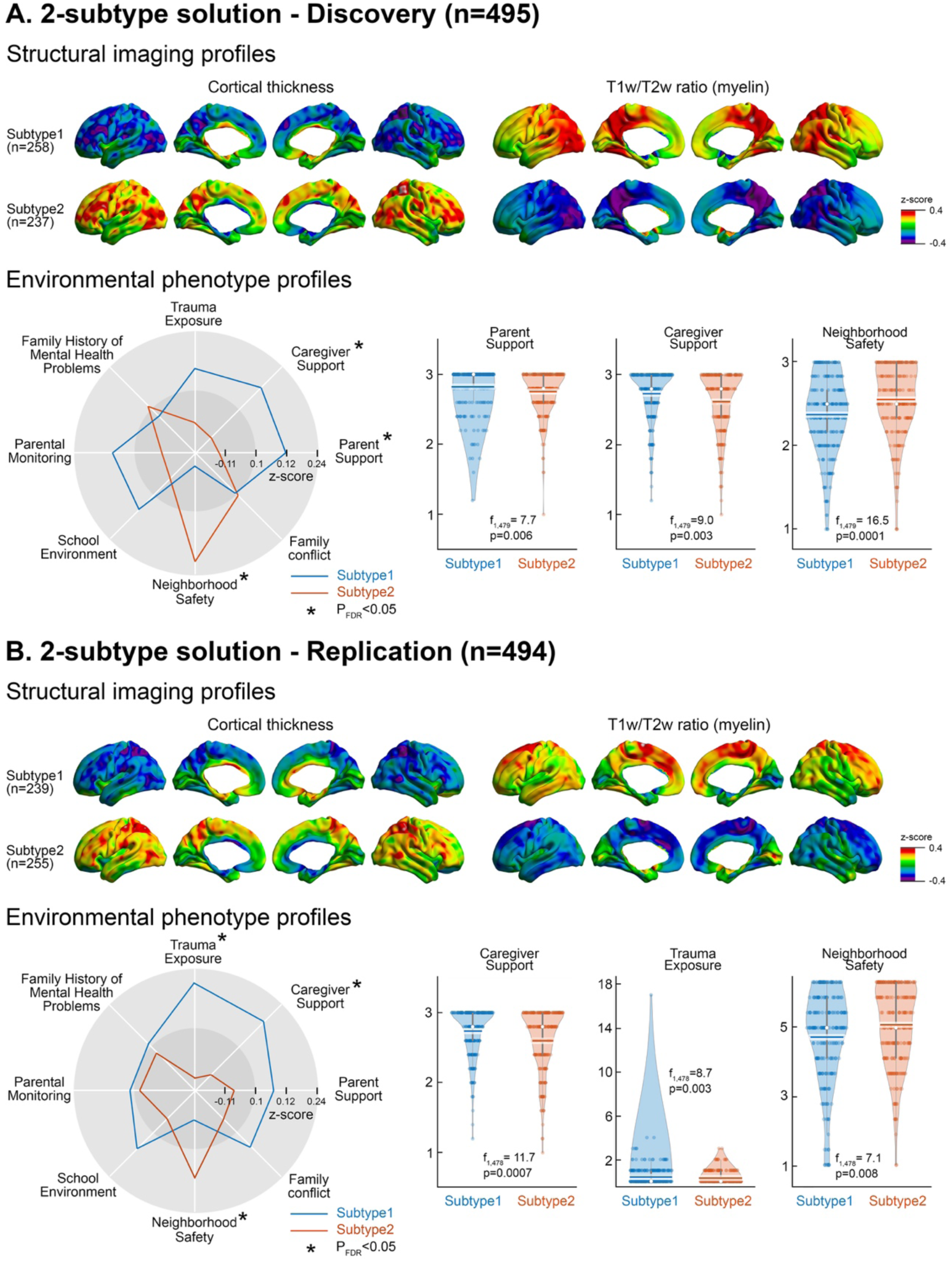
2-subtype solution – Discovery and Replication. The two-subtype solution is presented for both discovery (A) and replication (B) datasets. Profiles of cortical thickness and T1w/T2w ratio measures are shown for each subtype. The spider plots display multidimensional profiles (z-score) of environmental factors for each subtype, with an asterisk indicating a significant difference between the subtypes. The raw data for these significant differences in environmental factors are shown in the violin plots.

**Figure 3.**
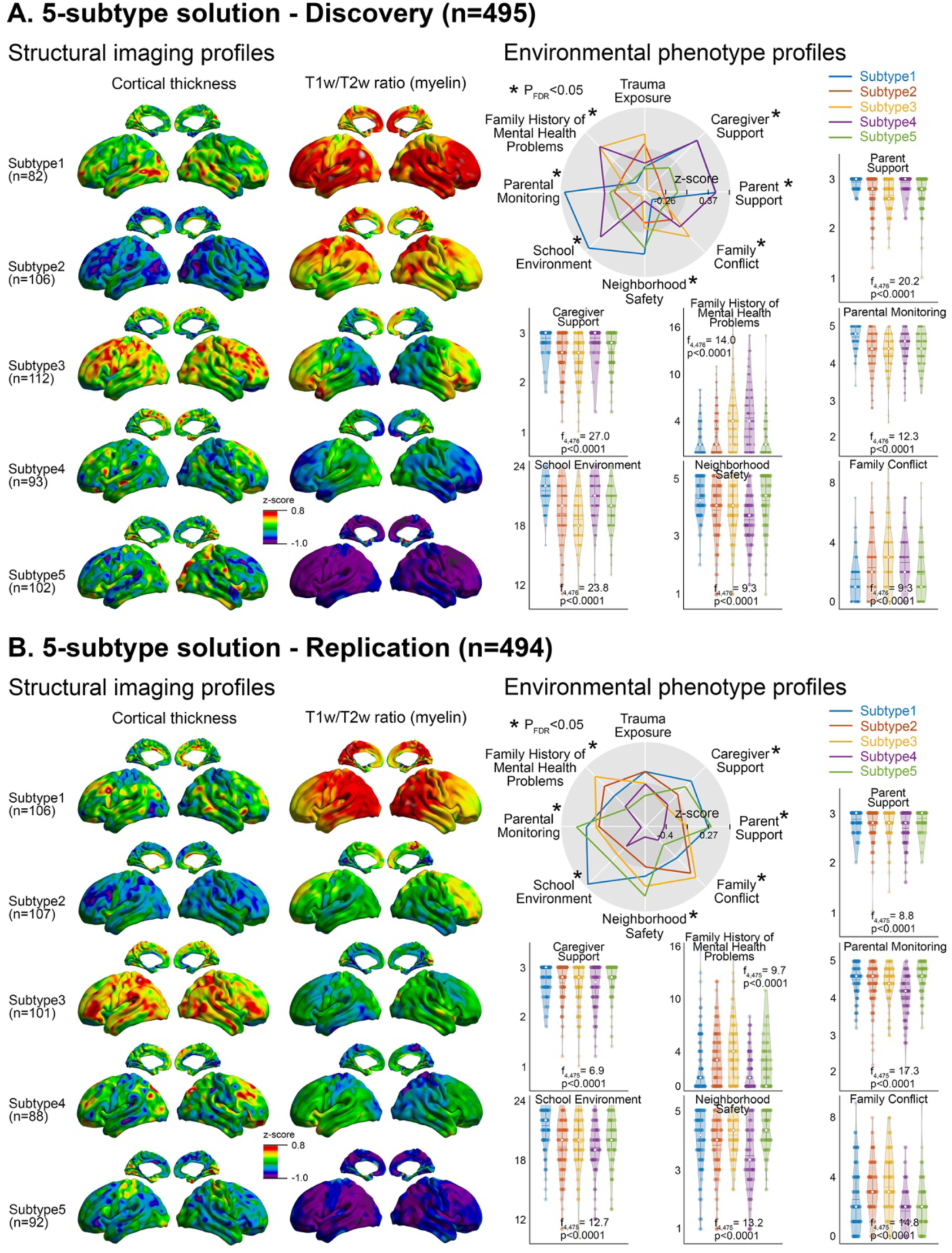
5-subtype solution – Discovery and Replication. The five-subtype solution is presented for both discovery (A) and replication (B) datasets. Profiles of cortical thickness and T1w/T2w ratio measures are shown for each subtype. The spider plots display multidimensional profiles (z-score) of environmental factors for each subtype, with an asterisk indicating a significant ANCOVA effect between the subtypes. The raw data for these significant differences in environmental factors are shown in the violin plots.

#### i) Two-subtype solution

The number of participants classified into each of these two subtypes was balanced (Subtype 1 n=258, Subtype 2 n=237), suggesting that their subtype-specific brain imaging and environmental phenotypic profiles were equally well represented across the sample. The brain imaging profiles revealed inverse patterns between cortical thickness and myelin across the two subtypes (**Figure 2** ‘*structural imaging profile*’). Indeed, Subtype 1 displayed slightly reduced cortical thickness across the whole brain (z-score: −0.4 to −0.2), and increased myelin (z-score: 0.20-0.35), particularly in the posterior cortical areas (relative to Subtype 2). Subtype 2 displayed opposite features of increased cortical thickness and reduced myelination (relative to Subtype 1). An ANCOVA analysis (in this case, equivalent to a two-sample t-test, given there were two subtypes) confirmed these subtype-specific characteristics at both the global and regional level (FIGURE 4A). Specifically, both cortical thickness and T1w/T2w ratio measures showed marked global differences in the whole-brain mean (discovery/replication: decrease in Subtype 1 *vs*. increase in Subtype 2, F_1, 478_=430/660, p<0.0001 for cortical thickness; increase in Subtype 1 *vs*. decrease in Subtype 2, F_1, 478_=52/10.2, p<0.0002 for T1w/T2w ratio). The subsequent ANCOVA targeting regional differences between the subtypes (*i*.*e*., ANCOVA statistically correcting for whole-brain mean) also showed such differential patterns, with two overlapping clusters that were significant in both the discovery and replication data. Indeed, the T1w/T2w ratio showed overlapping clusters (*i*.*e*., reproducible subtype differences) in the right paracentral lobule and left superior frontal cortices (p_RFT_<0.05). There were no regional differences between the subtypes in cortical thickness.

**Figure 4.**
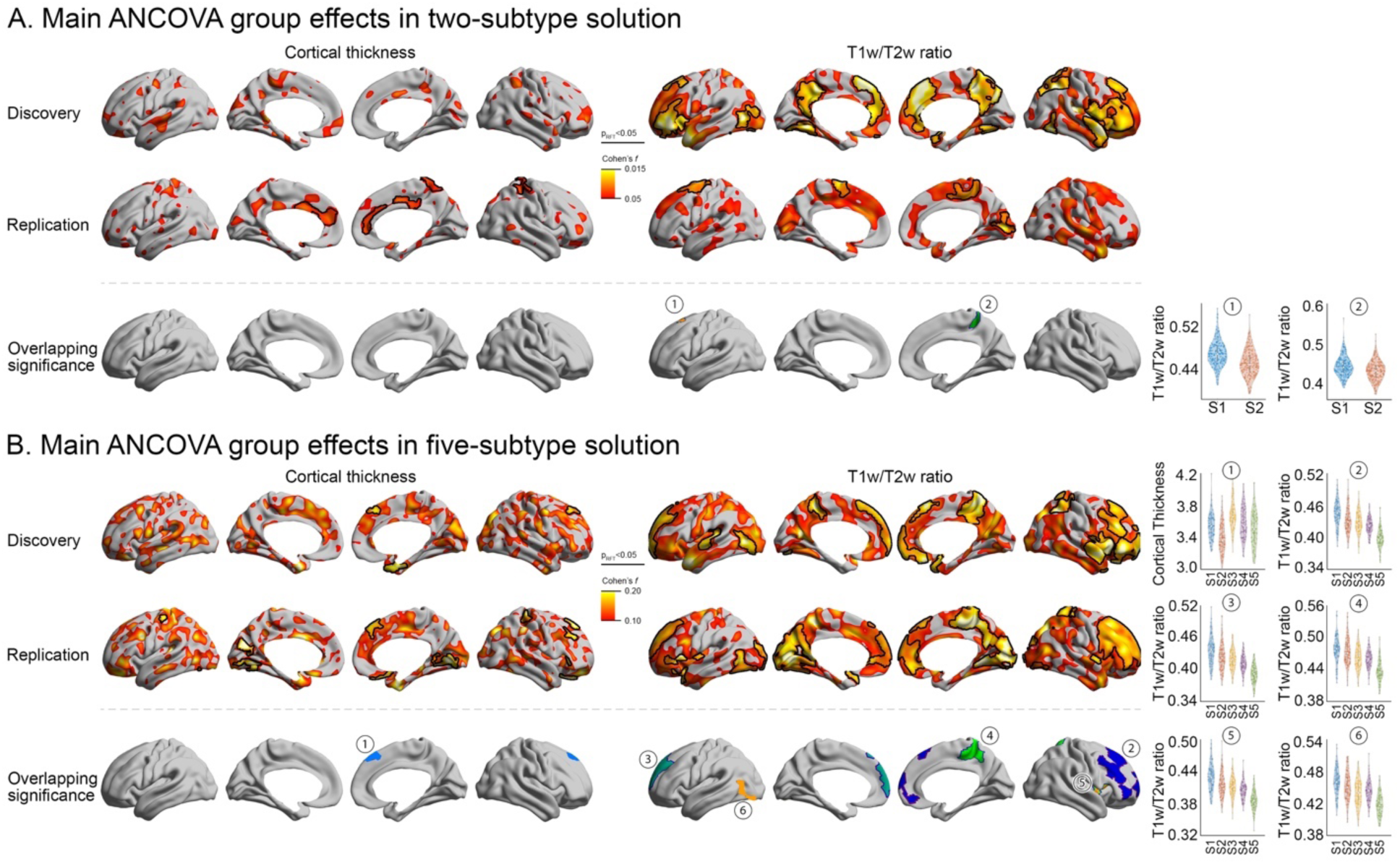
Overlapping subtype differences across discovery and replication datasets. ANCOVA was performed to evaluate group differences in cortical thickness (left) and T1w/T2w ratio (right) for the two-subtype (**A**) and five-subtype (**B**) solutions (See ‘*Subtype profiling*’ in **Methods and Materials** for statistical details). The main group effect was displayed by a Cohen’s *f* squared value 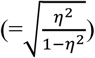 (Lakens, 2013), with black boundaries indicating significant clusters based on random field theory (RFT; p_RFT_<0.05). Brain areas with effects that survived RFT in both the discovery and replication results are presented (‘Overlapping significance’) to highlight the reproducible main group effects. For each overlapping cluster, the distribution of brain imaging features across individuals is shown for the identified subtypes at the right.

Individuals in Subtype 1 versus Subtype 2 differed in the extent to which their environments were characterized by parental support, caregiver support, and neighborhood safety (ANCOVA: FDR<0.05; **Figure 2** ‘*Environmental phenotype profile*’), but there were no differences in trauma exposure, family history of mental health problems, parental monitoring, or school engagement. Specifically, individuals in Subtype 1, characterized by reduced cortical thickness and increased myelination, had higher support (*e*.*g*., positive evaluation and affection) displayed by parents and caregivers and lower perceptions of neighborhood safety than individuals in Subtype 2. Subtypes 1 and 2 did not significantly differ in CBCL T-scores for internalizing symptoms, externalizing symptoms, or total problems (uncorrected p>0.05).

#### ii) Five-subtype solution

Similar to the two-subtype solution, the number of participants classified in a given subtype was also relatively balanced across the identified subtypes (Subtype 1: 82, Subtype 2 n=106, Subtype 3 n=111, Subtype 4 n=93, Subtype 5 n=103). However, there was greater variability across their brain imaging and environmental phenotypic profiles, suggesting that a higher number of subtyping solutions might present a more detailed picture of complex relationships between environmental factors and brain structure (**Figure 3**). Specifically, Subtype 1 showed more highly myelinated whole-brain patterns and a slight increase in cortical thickness in lateral temporo-occipital regions. Subtype 5 presented largely opposite patterns in both imaging features, demonstrating markedly decreased whole-brain myelin and mild cortical thinning in the right lateral frontal area. Subtypes 2-4 showed gradual changes in myelin patterns from high (Subtype 1) to low whole-brain T1w/T2w z-scores (Subtype 5) across multiple brain areas (**Figure 3** ‘*Structural imaging profile*’). The cortical thickness patterns also varied across the subtypes, showing widespread decreased thickness in Subtype 2, relative to increased thickness in Subtype 3, and an intermediate pattern in Subtype 4.

This qualitative subtype profiling was complemented by the following statistical analysis (ANCOVA), where we demonstrated reproducible group effects across identified subtypes. Compared to the two-subtype solution, these higher-resolution subtyping results showed more overlap in findings between the discovery and replication datasets. First, the global mean in both cortical thickness and T1w/T2w ratio measures demonstrated clear subtype-specific patterns that were highly reproducible (discovery/replication: F_4, 474_=430/186, p<0.0001 for cortical thickness; F_4, 474_=81.4/10.2, p<0.0002 for T1w/T2w ratio). Moreover, regional effects were also consistent across the two datasets for both cortical thickness and T1w/T2w ratio (**Figure 4B**). Subtype differences in cortical thickness were localized to the superior frontal area, whereas differences in myelin were primarily located in the default mode and frontoparietal networks (p_RFT_<0.05). These findings suggest that brain-based changes related to heterogeneous environmental conditions may not stem from a single overarching developmental process, but possibly from both anatomically-nonspecific global changes and regionally-specific modulation.

Environmental profiles also varied across the subtypes for parent and caregiver support, family conflict, neighborhood safety, and school environment (ANCOVA: FDR<0.05) (**Figure 3** ‘*Environmental phenotype profile*’). However, there were no differences in trauma exposure between the subtypes. Similar to observed patterns in the two-subtype solution (see *Figure 2*), Subtype 1 was characterized by higher myelination and increased parental and caregiver support. Although the opposite association was seen in Subtype 4 (*i*.*e*., reduced myelination and increased parental and caregiver support; **Figure 3** for discovery data), the degree of reduction in myelination was slight, and this group also had more severe family conditions (*e*.*g*., family mental health problems and criminal history) and lower parental monitoring than Subtype 1. More importantly, the Subtype 4 patterns, especially for parent and caregiver support, were not clearly observed in the replication dataset. Another notable association between environmental and brain features was observed in Subtype 3. Specifically, this subgroup was characterized by less favorable conditions in almost every domain of environmental factors (*i*.*e*., lower parental and caregiver support, lower parental monitoring, a more significant family history of mental health problems, less favorable school environment, and higher family conflict) and relative cortical thickening without alterations in myelination. Importantly, the majority of these findings in both the two- and five-subtype solutions were reproduced in the independent replication data, confirming their high generalizability.

Finally, we examined differences in clinical symptom scores between subtypes using a general linear model, controlling for age and sex, and correcting for site of acquisition (see **Supplementary Figure 3**). Given the number of pairwise tests conducted, we only report results that survived FDR comparison. For internalizing symptoms, Subtype 3 was observed to have higher scores than Subtype 1 (p=0.012). For externalizing symptoms, Subtype 3 had higher scores than subtypes 1 (p<0.0001), 4 (p=0.0027), and 5 (p=0.007), and Subtype 2 had higher scores than Subtype 1 (p=0.0044). With regard to total problem scores, Subtype 3 again had higher scores than Subtype 1 (p=0.002), Subtype 4 (p=0.0114), and Subtype 5 (p=0.0018). These patterns largely replicated in the independent dataset, with Subtype 3 having higher scores across internalizing symptoms, externalizing symptoms, and total problems than other subtypes (p<0.001).

Of note, SNF findings based on either brain imaging or phenotypic scores alone did not yield comparable subtype profiles. When subtyping based on only brain imaging data, the imaging features revealed similar structural profiles across the identified subtypes compared to our main findings (*e*.*g*., opposite cortical thickness and myelin patterns between the two subtypes). However, there were no between-subtype differences on environmental factors, and we were unable to predict clinical scores based on subtypes derived from brain imaging data alone. When subtyping based on only environmental data, SNF detected statistically more robust subtype-specific environmental profiles. However, brain structural features did not differ across the subtypes. These findings suggest that the SNF approach considering multimodal data effectively extracted unique brain-environment relationships, which could not be captured by unimodal approaches.

### Subtype validation

To validate the significance of identified subtypes, we performed two independent prediction analyses. The first analysis assessed how generalizable the brain structural and environmental phenotypic profiles for each subtype were in unseen cases through subtype classification. The second analysis validated the subtypes by using the brain imaging features to predict clinical symptom scores in unseen cases.

#### i) Prediction of subtype membership classification (**Figure 5A**)

For the two-subtype solution, the prediction result based on the replication data showed correct subtype classification in 86% of the cases (chance level estimated by a permutation test: 48%; p<0.001), suggesting that the brain structural and environmental profiles learned from the discovery dataset for each subtype were highly generalizable. Notably, mapping the weight of the features learned by a support vector machine revealed that regions of the sensorimotor and default networks had particularly high influence in distinguishing the two subtypes. Although classification accuracy was lower, generalizable subtype-dependent data profiles were also found in the five-subtype solution, which revealed successful classification in 63% of the cases (chance level=19%; p<0.001). The feature contribution map highlighted similar brain areas (i.e., sensorimotor and default networks) as found in the two-subtype classification, which suggests that these anatomical regions may be particularly sensitive to the environment during childhood.

**Figure 5.**
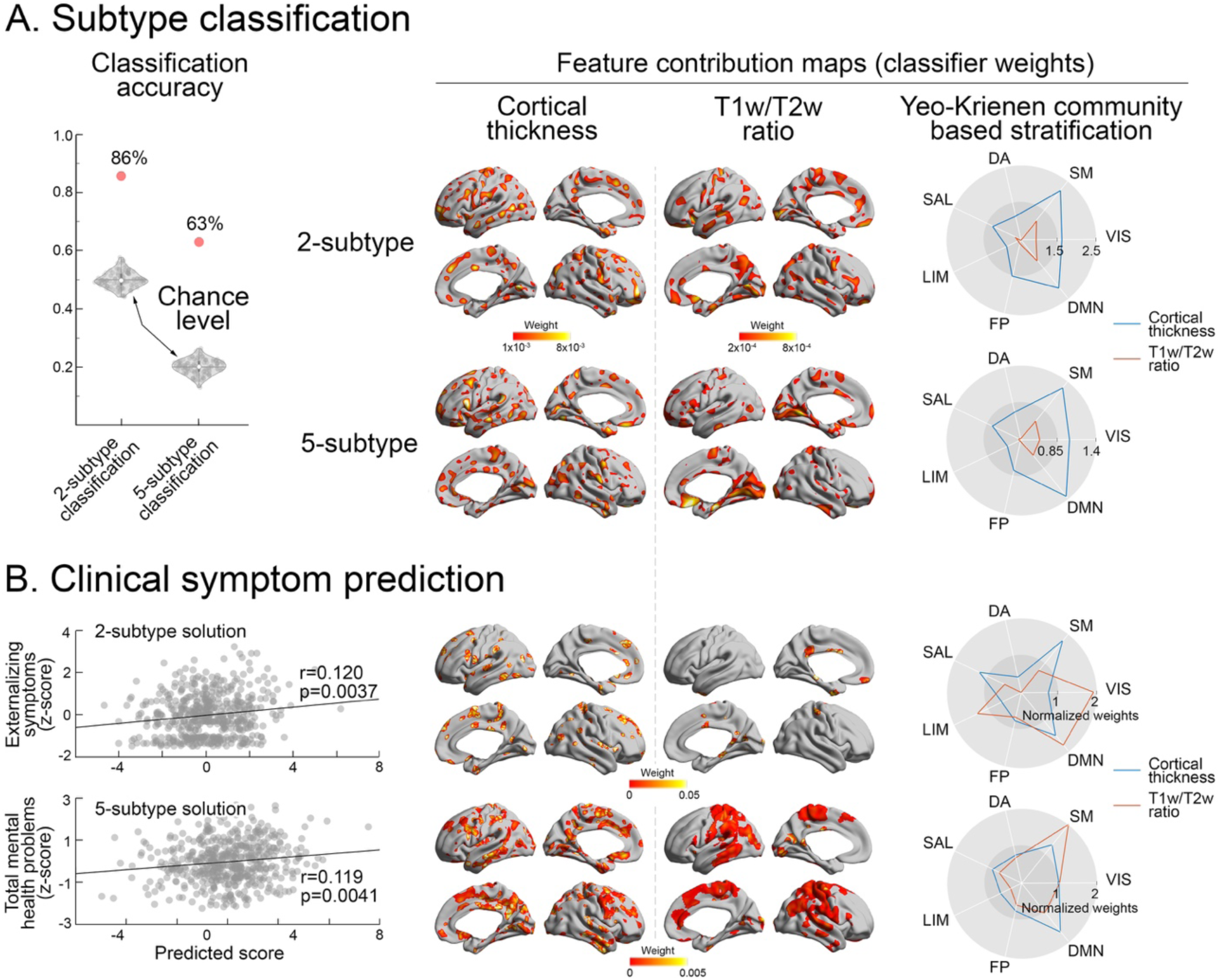
Subtype classification and clinical symptom prediction. Results are shown for the prediction of subtype classification (A) and clinical symptoms (B). A) The red dots in the violin plot represent classification accuracy in the 2- and 5-subtype solutions. To demonstrate their significance against a chance level, we performed permutation tests 100 times, randomly shuffling the subtype membership in the training (discovery) dataset. B) The correlations between actual clinical scores (upper: externalizing symptoms, lower: total mental health problems) and predicted scores are displayed in the scatter plot. In both (A) and (B), the feature contributions were identified based on the weights provided from the classifiers and mapped on the whole brain and stratified using the Yeo-Krienen functional community atlas (B. T. T. Yeo et al., 2011). Prediction accuracy for externalizing symptoms was informed by the 2-subtype solution; prediction accuracy for total mental health problems was informed by the 5-subtype solution. Abbreviations: VIS=visual, SM=somatosensory, DA=dorsal attention, SAL=salience, LIM=limbic, FP=frontoparietal, DMN=default mode network.

#### ii) Prediction of clinical symptoms (**Figure 5B**)

The prediction of clinical scores based on support vector regression showed marginal but significant correlation between actual and predicted scores. Specifically, the classifier informed by the two-subtype solution successfully predicted externalizing symptoms (Spearman correlation: r=0.120, p=0.0037; survived FDR comparison) but not internalizing symptoms or total problems (r=-0.08/0.06, p=0.96/0.086, respectively). The classifier informed by the five-subtype solution successfully predicted total problem scores (r=0.119, p=0.0041; survived FDR comparison), but not internalizing or externalizing symptoms (r=0.07/0.08, p=0.054/0.03, respectively; did not survive FDR comparison). The features contributing to the significant prediction of externalizing problems were mainly found in the primary sensory and higher-order default mode systems. Notably, the feature map for the prediction of total problems demonstrated similar contribution profiles, implicating both lower-level somatosensory and higher-order default mode areas. By contrast, when we performed this prediction across all participants without subtype information, we failed to predict any clinical symptom scores.

To assess the robustness of our results, we iteratively performed the same prediction analysis 100 times across bootstrapped samples (resampling 90% of cases without replacement). We found that indeed the majority of prediction results centered around the original prediction conducted on all participants, suggesting that this performance was not driven by outliers. Another noteworthy finding in this bootstrap analysis was that even for the non-significant prediction results (*e*.*g*., total problems in the two-subtype solution, externalizing symptoms in the five-subtype solution), their distributions showed higher prediction accuracy compared to that of the subtype-free approach (p<0.001; **Supplementary Figure 2**). These results collectively emphasize that given a highly heterogeneous sample, identifying more homogeneous subgroups may be an optimal strategy before performing brain-phenotype association analyses.

Overall, these analyses demonstrate that the observed subtypes are highly reproducible, and that subtype classification aids in decomposing heterogeneity in brain structure and environmental exposures to enhance efficacy of clinical prediction. Moreover, the sensorimotor and default mode networks found to be influential for both subtype classification and clinical prediction may represent developmentally sensitive brain systems. Such systems may be readily affected by the childhood environment and thus related to the manifestation of mental health in school-aged children.

## DISCUSSION

In this study we dissected the complex relationships between the childhood environment and brain structure using a fully unsupervised, multimodal data integration and clustering approach in a large sample of school-aged children from the ABCD Study. The method used here, Similarity Network Fusion (SNF), combined two well-established developmental MRI features, namely cortical thickness and myelin-surrogate markers, and key environmental risk and protective factors to identify distinct subtypes. This approach identified two viable solutions, one with two subgroups and one with five (depending on the clustering resolution), each showing divergent patterns of brain structure and environmental exposure. Across both solutions, patterns of brain cortical thickness and myelination were highly replicable, with consistent global and regional differences in thickness and myelination observed between subtypes. Patterns of co-occurrence between specific environmental exposures and neural phenotypes varied across the subtypes, revealing particularly strong associations between brain structure and supportive caregiving and neighborhood safety. Importantly, by leveraging subtype classification information, we were able to predict externalizing symptoms and total mental health problems in independent unseen cases (replication data), which was not possible without subtype classifications. Finally, the lack of meaningful subtyping based on either brain structure or environmental factors alone highlights the value of integrating across modalities to better represent the relationships between neural and environmental factors. In sum, the present study provides a proof-of-concept demonstration of the value of a multimodal and fully data-driven approach to decompose heterogeneous environmental effects on brain development.

Findings from the two-subtype solution provided novel insight into overarching associations among brain and environmental factors, while findings from the five-subtype solution further parsed heterogeneity to produce relatively more homogenous clusters of environmental exposures and brain structure. Overall, the two-subtype solution demonstrated reciprocal patterns across subtypes. In one subtype, global cortical thinning and increased myelination co-occurred with higher levels of parent and caregiver support as well as decreased neighborhood safety. In the other subtype, the inverse pattern occurred, with increased global cortical thickening, decreased myelination, decreased parent and caregiver support, and increased neighborhood safety. These findings were mostly replicated in an independent dataset, though parent support did not differ between the subtypes in the replication dataset. Caregiving was also central to the strongest findings in the five-subtype solution, where less supportive caregiving and higher family conflict co-occurred with globally increased cortical thickness (Subtype 3). These findings are generally consistent with evidence linking favorable caregiving conditions with increased cortical thinning. Within the five-subtype solution, differences in cortical thickness were particularly evident in the right superior frontal region. Prior research has shown that a higher degree of positive maternal behavior is associated with increased cortical thinning in the right anterior cingulate and bilateral orbitofrontal cortices among males (Whittle et al., 2014), whereas higher maternal aggressiveness has been linked with increased cortical thickening in the right superior frontal and lateral parietal cortices (Whittle et al., 2016). Further, supportive caregiving has been shown to buffer the effects of adversity related to poverty and living in a socioeconomically-disadvantaged neighborhood (Brody et al., 2019; Whittle et al., 2017). Specifically, male adolescents who had positive caregivers demonstrated relatively increased cortical thinning in dorsofrontal and orbitofrontal cortices compared to those who did not have supportive caregivers and were also living in disadvantaged neighborhoods (Whittle et al., 2017). Exposure to community violence has been associated with decreased gray matter volumes in adolescence (Butler et al., 2018; Saxbe et al., 2018), but less is known about perceptions of neighborhood safety and brain structure. Our findings extend previous work to demonstrate that the neural correlates of caregiving and neighborhood safety may go beyond changes in specific brain regions and involve global shifts in cortical thickness.

Though less is known about myelination and childhood experiences, the co-occurrence of supportive caregiving and increased myelination across both the two- and five-subtype solutions may be consistent with normative development, as myelination increases throughout development (Lebel & Deoni, 2018). Regional differences in myelination between the five subtypes were observable in clusters located in the right medial and lateral frontal convexity, the posterior cingulate cortex and paracentral lobule, the lateral occipito-temporal area, and the insula. Within the five-subtype solution, despite only moderate cortical thinning, the highest levels of myelination co-occurred not only with high caregiving support but also with high parental monitoring, lower family conflict, a lesser family history of mental health problems, higher neighborhood safety, and a more positive school environment (Subtype 1). These patterns provide novel evidence that myelination and cortical thickness may be independently associated with discrete environmental exposures. A recent study indicated that cortical thinning throughout child development largely results from increasing myelination. Yet changes in myelination did not fully explain variability in cortical thinning, suggesting that various cellular processes also contribute to cortical morphology (Natu et al., 2019). Our findings appear to align with this model. Specifically, inverse associations between myelination and cortical thickness were observable across the two- and five-subtype solutions. However, unique associations between specific environmental exposures and myelination versus cortical thickness in the five-subtype solution suggest that environmental exposures may differentially impact these processes.

Given the demonstrated relationships between childhood experiences and psychopathology, and the potential for changes in brain structure to contribute to this effect, we examined the clinical relevance of these subtypes. Despite robust differences in neural and environmental phenotypes, there were no differences in clinical symptoms in the two-subtype solution. However, symptoms did differ between subtypes in the five-subtype solution, consistent with the idea that this solution may have parsed heterogeneity more finely to produce meaningful groupings of brain and environmental factors, and their associations with clinical symptoms. Most notably, Subtype 3 displayed the highest internalizing, externalizing and total problems across both the discovery and replication datasets. The profile of this subtype, characterized by family conflict and history of psychopathology and globally increased cortical thickness, aligns with previous work linking internalizing and externalizing symptoms with reduced cortical thinning in late childhood (Whittle et al., 2020), and externalizing problems with attenuated cortical thinning in adolescence (Oostermeijer et al., 2016). By contrast, other studies have found reduced cortical thickness in association with externalizing (Ameis et al., 2014; Ducharme et al., 2011) and internalizing (Bora et al., 2012; Newman et al., 2016) disorders, highlighting the importance of future work to address the challenging nature of linking brain, environment, and clinical outcomes. Here we also found that subtyping enhanced clinical prediction, such that including subtype classification information enhanced the prediction of clinical symptoms based on neuroanatomical features (which was not possible without the inclusion of subtype information). These findings suggest that subtyping approaches may prove useful for understanding heterogeneity in clinical outcomes during development.

This study leveraged the data from the ABCD Study, a large publicly available and demographically diverse, population-based developmental cohort, to test whether heterogeneity in brain structure and environmental exposures could be parsed using a multimodal data fusion and subtyping approach. To our knowledge, this study is the first to examine associations between myelination and cortical thickness with environmental risk and protective factors across multiple levels (*i*.*e*., caregiver, family, school, neighborhood). A key strength of this work is the dimensional approach taken to assess environmental exposures. Given that much previous work has focused on solely adverse experiences, or examined only one facet of the environment (*e*.*g*., caregiving), we prioritized using a data-driven method to empirically determine patterns of co-occurrence between different types of environmental exposures and measures of brain structure. Our findings demonstrated highly reproducible and robust brain-environment relationships, and the confirmation of our results in a held-out sample is an important strength of this work. Finally, we applied the subtyping of brain structure and environmental exposures to examine individual differences in clinical symptoms. As this study employed the first wave of ABCD data, the findings set the stage for future investigations of how adversity and brain structure are associated throughout adolescence, as well as how they may predict longitudinal changes in mental health. This vast potential for follow-up research is well aligned with the ABCD Study’s vision of open science and collaborative research to facilitate novel insight into brain and behavioral development.

While this study meaningfully contributes to the literature on childhood experiences and brain development, there are nevertheless aspects upon which future research can improve. Here we employed a population-based dataset derived from communities nationwide and tested generalizability in unseen cases within this same sample. However, replication of these findings in external samples will be needed to confirm generalizability. The age range of 9-10 years old in this sample also represents a very specific point in childhood, and it is unknown whether similar brain-environment relationships would be observed at different ages during development. Given marked neurodevelopmental changes throughout childhood and adolescence (Casey et al., 2019; Gee et al., 2018; Giedd et al., 1999; Herting & Sowell, 2017; Kaczkurkin et al., 2019; Luna, 2009), as well as the likelihood that different environmental factors will influence the brain in unique ways depending on developmental stage (Cohodes et al., 2020; Gee & Casey, 2015; Lupien et al., 2009; Tottenham & Sheridan, 2010), conducting subtyping of brain-environment relationships across development is a vital next step. Perhaps due to the relatively low rate of parent-reported trauma exposure in the current sample, childhood trauma did not meaningfully vary across subtypes in this study. Moreover, we were unable to examine differential effects of specific types of trauma given the limited variability in reported exposures. Given the well-documented link between childhood trauma and psychopathology (Green et al., 2010), examining trauma at later ages and with child self-report will be important next steps. The present study observed notable differences in cortical thickness and myelination between subtypes but only examined these two features of brain structure. In order to gain a more nuanced understanding of differential patterns of brain structure, future research will benefit from assessing additional measures such as white matter integrity or developmentally relevant morphological features (*e*.*g*., sulco-gyral profiles). Evaluating the clinical relevance of the current findings is another important goal for future work. Though our findings demonstrate that the current subtypes inform prediction of contemporaneous externalizing symptoms, testing prediction of future symptoms and disorder onset will be essential to assessing the clinical utility of these subtypes. Similarly, longitudinal data will be important for examining the temporal nature of changes in brain structure and externalizing symptoms across development, which was not possible within the current cross-sectional study. Finally, due to the already high-dimensional nature of the data, we limited our current investigation to brain structure, environmental factors, and clinical symptoms. However, future examinations employing similar approaches could benefit from integrating the functional neuroimaging data and genetic data available in the ABCD Study to provide a deeper understanding of the biological processes related to the childhood environment and mental health outcomes.

Taken together, the current study supports the utility of subtyping approaches for examining associations between brain structure and environmental exposures during development. Among the observed brain-environment patterns, findings highlighted the co-occurrence of supportive caregiving and lower family conflict with increased cortical thinning. These patterns underscore the importance of family functioning during middle childhood and point to caregiving as a potential target for interventions designed to bolster resilience in the face of adversity. Finally, this study demonstrates that parsing heterogeneity in brain and environmental exposures has clinical relevance, and provides a foundation on which future studies can leverage subtyping approaches to predict mental health outcomes across development.

## ACKNOWLEDGEMENTS

This work was supported by funding from the National Institutes of Health (NIH) Director’s Early Independence Award (DP5OD021370), National Institute on Drug Abuse (U01DA041174), Brain & Behavior Research Foundation (NARSAD Young Investigator Award), Jacobs Foundation Early Career Research Fellowship, and The Society for Clinical Child and Adolescent Psychology (Division 53 of the American Psychological Association) Richard “Dick” Abidin Early Career Award and Grant for DGG; the Brain & Behavior Research Foundation (NARSAD Young Investigator Award #28436), the Canadian Institutes of Health Research (postdoctoral fellowship MFE-158228), and the Institute for Basic Science (IBS-R15-D1) for SJH; and the National Institute of Mental Health (U01MH099059) and an endowment from Phyllis Green and Randolph Cowen for MPM.

## SUPPLEMENTARY MATERIAL

**Supplementary Table 1.** Inclusion/exclusion criteria and quality control.

|  | Participants<br>remaining | Sites<br>remaining | Scanners<br>remaining |
| --- | --- | --- | --- |
| ABCD Release 2.0 | 11500 | 22 | 5 |
| Participants with data of interest <sup>a</sup> | 5450 | 14 | 3 |
| Siblings excluded | 4448 | 14 | 3 |
| Participants with autism spectrum disorder or<br>epilepsy excluded | 4339 | 14 | 3 |
| Participants with affected sMRI images excluded <sup>b</sup> | 3721 | 14 | 3 |
| FreeSurfer QC passed <sup>c</sup> | 2379 | 14 | 3 |
| Final random sampling <sup>d</sup> | 1000 | 13 | 2 |

a. T1- and T2-weighted MRI, 8 environmental scores, 3 clinical scores.
b. n=1137 participants had their scans incorrectly flipped; n=1 incorrectly associated with a different pGUID (ABCD Study: Known issues with Data Release 2.0).
c. No presence of motion artifacts, pial mater overestimation, white matter underestimation, or field inhomogeneity.
d. 1000 participants randomly sampled.

**Supplementary Table 2.** Demographic profiles of 2-/5-subtype solutions (Replication)

| 2-subtype profiling | Subtype 1<br>(n=239) | Subtype 2<br>(n=255) | p value |
| --- | --- | --- | --- |
| Age (months; mean ±SD) <sup>a</sup> | 119.3±6.9 | 119.6±7.1 | p=0.65 |
| Sex (male/female) <sup>b</sup> | 124/115 | 130/125 | p=0.84 |
| Site (1<sup>st</sup>-13<sup>th</sup> sites in order) <sup>b</sup> | 15/11/12/24/8/14/24/7/15/21/<br>49/12/27 | 19/17/18/31/8/16/22/5/19/15/<br>44/12/29 | p=0.97 |
| Race <sup>b, c</sup> | 175W, 28B, 3A, 8O, 23M, 2NA | 178W, 30B, 5A, 7O, 33M, 2NA | p=0.86 |
| Ethnicity <sup>b, d</sup> | 44H, 193 nH, 2NA | 60H, 194nH, 1NA | p=0.17 |
| Education level of parent <sup>b, e</sup> | 40E/M/H, 148B, 51G | 24E/M/H, 154B, 77G | p=0.01 <sup>g</sup> |
| Household income <sup>b, f</sup> | 55L, 72M, 97H, 15NA | 57L, 73M, 113H, 12NA | p=0.78 |

| 5-subtype profiling | Subtype 1<br>(n=106) | Subtype 2<br>(n=107) | Subtype 3<br>(n=101) | Subtype 4<br>(n=88) | Subtype 5<br>(n=92) | p value |
| --- | --- | --- | --- | --- | --- | --- |
| Age (months; mean ±SD) <sup>a</sup> | 119.6±7.1 | 119.5±6.9 | 119.7±6.9 | 118.3±7.0 | 120.0±7.4 | p=0.56 |
| Sex (male/female) <sup>b</sup> | 57/49 | 51/56 | 51/50 | 42/46 | 53/39 | p=0.59 |
| Site (1<sup>st</sup>-13<sup>th</sup> sites in order) <sup>b</sup> | 7/6/5/13/4/9/<br>9/3/9/8/19/5/<br>9 | 7/9/5/12/3/6/<br>10/1/6/8/19/<br>7/14 | 6/3/13/6/2/5/<br>11/1/8/8/21/<br>4/13 | 6/3/2/12/6/2/<br>10/3/6/3/19/<br>5/11 | 8/7/5/12/1/<br>8/6/4/5/9/<br>15/3/9 | p=0.83 |
| Race <sup>b, c</sup> | 71W, 13B,<br>1A, 5O,<br>14M, 2NA | 80W, 13B,<br>1A, 1O,<br>12M | 73W, 11B,<br>2A, 2O,<br>13M | 62W, 10B,<br>2A, 5O, 8M,<br>1NA | 67W, 11B,<br>2A, 2O,<br>9M, 1NA | p=0.93 |
| Ethnicity <sup>b, d</sup> | 28H, 78nH | 18H, 87nH,<br>2NA | 20H, 81nH | 22H, 66nH | 16H, 75nH,<br>1NA | p=0.36 |
| Education level of parent <sup>b, e</sup> | 19E/M/H,<br>63B, 24G | 15E/M/H,<br>68B, 24G | 2E/M/H,<br>67B, 32G | 15E/M/H,<br>49B, 24G | 13E/M/H,<br>55B, 24G | p=0.04 <sup>g</sup> |
| Household income <sup>b, f</sup> | 24L, 32M,<br>42H, 8NA | 25L, 28M,<br>46H, 8NA | 23L, 30M,<br>43H, 5NA | 18L, 23M,<br>46H, 1NA | 22L, 32M,<br>33H, 5NA | p=0.63 |

a. Group comparison was based on independent samples t-test.
b. Group comparisons were based on Chi-square test (if all frequencies $\geq 5$ ) or Fisher's exact test (if any frequency $< 5$ ) of the contingency table.
c. W (White), B (Black), A (Asian), O (Other race), M (Mixed race), NA (Not answered or refused to answer); Details about race can be found here: https://nda.nih.gov/data_structure.html?short_name=pdem02.
d. H (Hispanic), nH (non-Hispanic), NA (Not answered or refused to answer).
e. E/M/H (Elementary/middle/high school), B (Bachelors), G (Graduate [Masters/PhD/Specialized degree such as MD]); Education information was based on the parent who completed the survey.
f. L(Low): $< \$50,000$ ; M(Middle): $\$50,000 \leq \text{income} < \$100,000$ ; H(High): $\leq \$100,000$ ; NA (Not answered or refused to answer). Household income was based on the sum of the past 12 months of gross pay for both parents.
g. Only uncorrected significance.

**Supplementary Figure 1.**
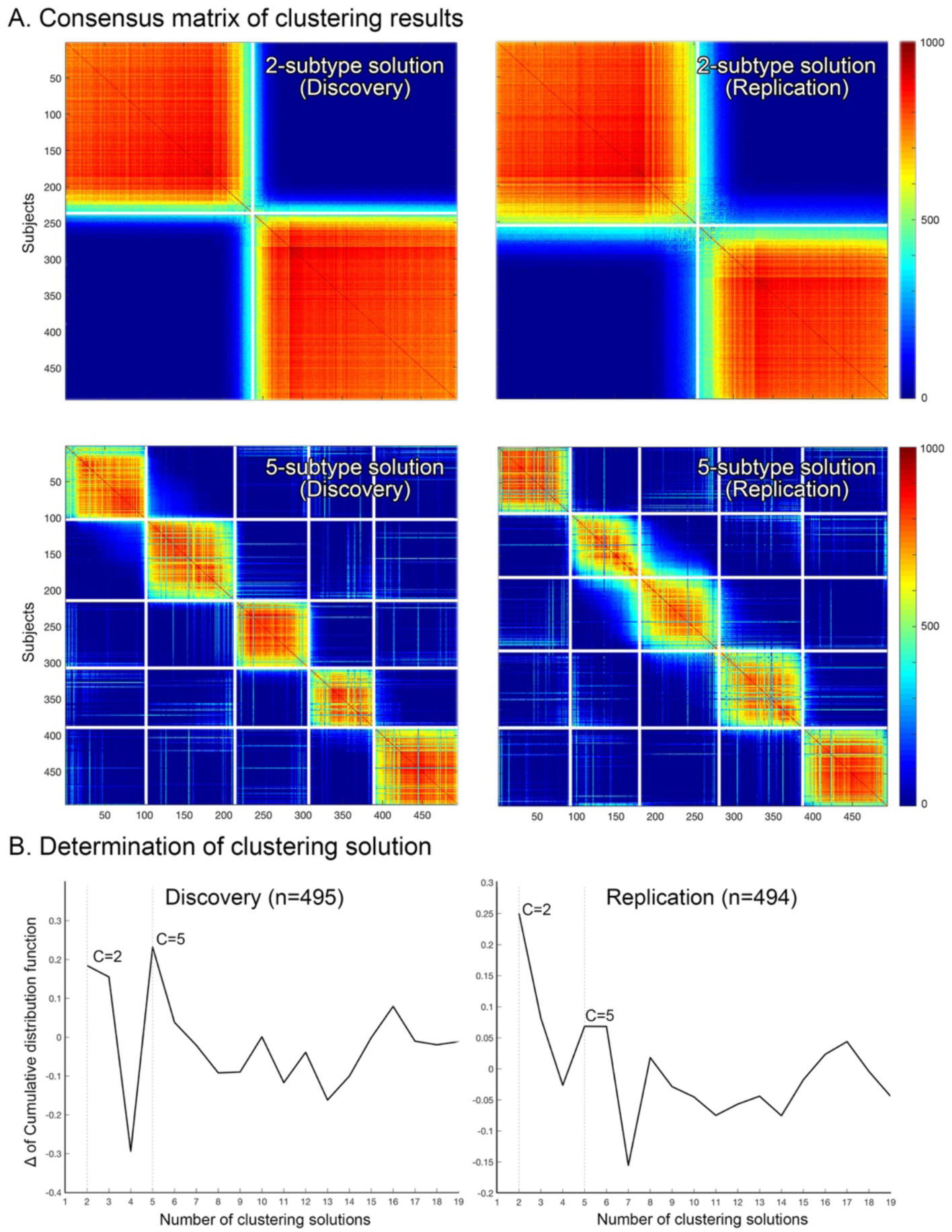
Finding a clustering solution based on consensus matrices. (A) 1000 iterations of bootstrapping (random sampling of 90% of participants) constructed a consensus matrix for a spectral clustering solution, which indicates how many times a pair of participants is clustered into the same group among 1000 iterations. (B) The degree of consensus at a given cluster number *C* is calculated as follows:

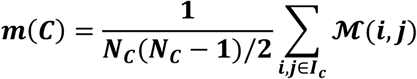

where *I*_*c*_ is the set of indices of items belonging to cluster *C*, that is, *I*_*c*._ = {j: *e*_c_ ∈ *C*}, N_c_ is the number of items in cluster *C*, and *M*(*I, j*) is the value of a consensus matrix at i^th^ row and j^th^ column. After calculating the degree of consensus m(*C*), a cumulative distribution function (CDF) can be estimated. The difference (Δ) of CDF between *C* and *C*-1 represents how much the consensus level can be increased by changing the clustering number from *C*-1 to *C*. We assessed which *C* produced the highest value in both the discovery and replication datasets, which resulted in *C*=2 and *C*=5. Their consensus matrices are shown in (A) as examples.

**Supplementary Figure 2.**
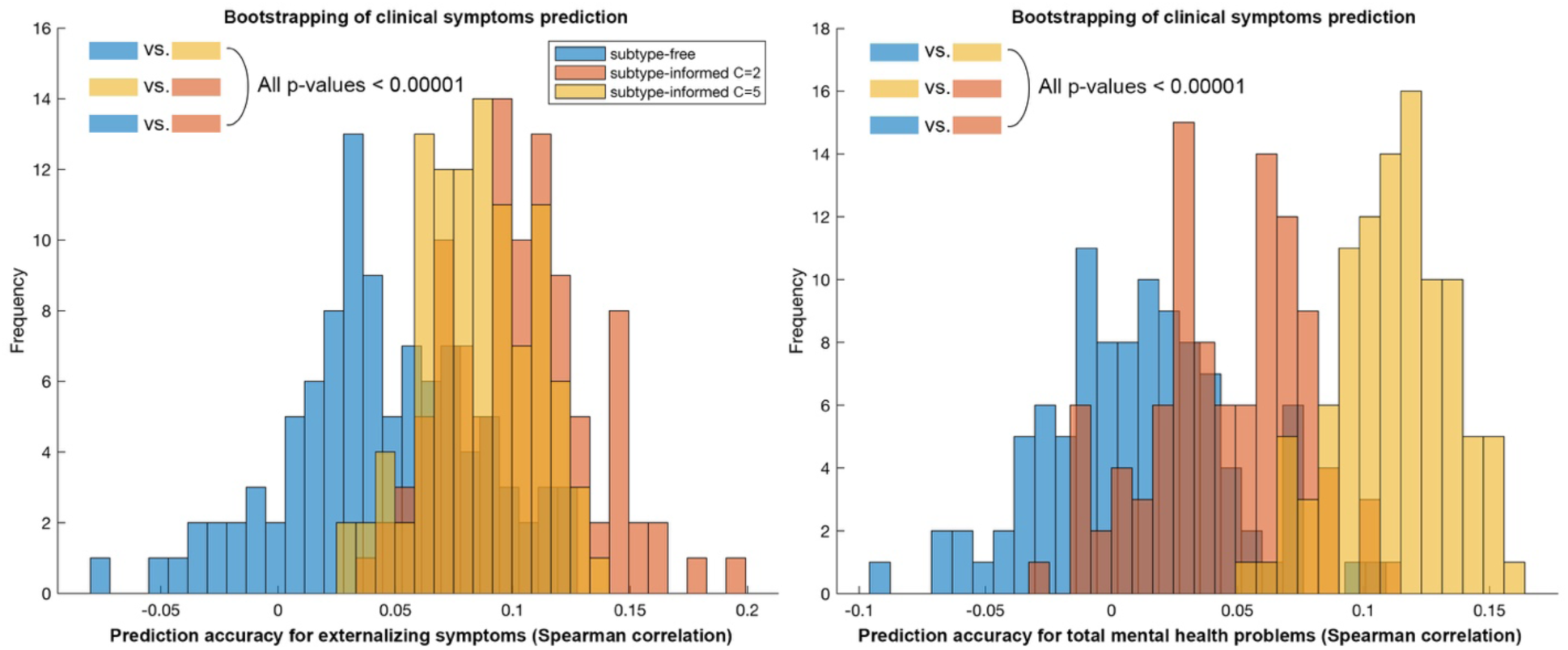
Bootstrap-based prediction analyses for clinical symptoms. To demonstrate generalizability of our main prediction results derived from all participants, we randomly selected 90% of participants, performed subtype-informed prediction for the three clinical scores (internalizing symptoms, externalizing symptoms, and total mental health problems), and repeated this analysis 100 times. Here we present the histogram of prediction accuracy from the clinical symptoms (*i*.*e*., externalizing symptoms and total mental health problems) showing the significance (after FDR correction) in the main analysis, which are the predictions of externalizing symptom (*left*, based on the 2-subtype solution) and total mental health problems (*right*, based on the 5- subtype solution) (see **Results** in the main text). Although prediction analyses based on the other subtype solution did not reach significance (yellow [five-subtype] in the left histogram and pink [two-subtype] in the right histogram), they still showed higher prediction accuracy compared to a subtype-free approach (blue), suggesting that the subtyping (*i*.*e*., finding more homogeneous subgroups) can improve phenotypic prediction. Indeed, there were significant differences (t statistics>4) in prediction accuracy (from the above histograms) between all pairs of conditions (*i*.*e*., subtype-free, two-subtype and five-subtype).

**Supplementary Figure 3.**
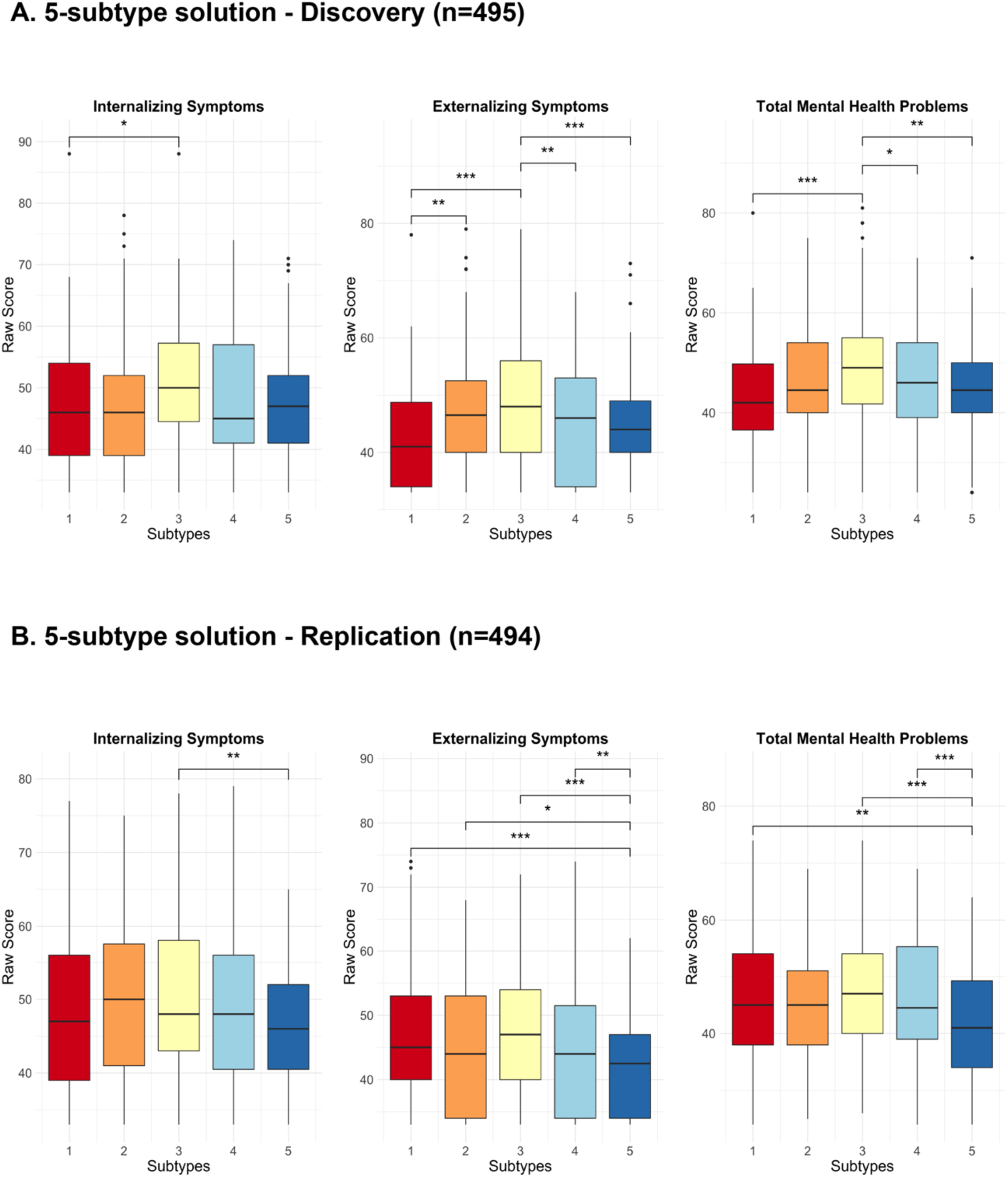
Group Differences in clinical symptoms in the 5-subtype solution. Using a generalized linear model, we conducted pairwise comparisons between each of the five subtypes, accounting for age and sex and controlling for site. The results shown here survived FDR correction (Benjamini & Hochberg, 1995).

## Environmental Factors

### Neighborhood Safety

The ‘Neighborhood Safety’ variable was computed from the ABCD Parent Neighborhood Safety/Crime Survey Modified from PhenX (NSC). We obtained the mean score across the three items listed below from the ABCD Sum Scores Culture & Environment measure (parent report).

| Item | Scoring |
| --- | --- |
| I feel safe walking in my neighborhood, day or night. | 1=Strongly Disagree, 2=Disagree, 3=Neutral, 4 = Agree, 5 = Strongly Agree |
| Violence is not a problem in my neighborhood. |  |
| My neighborhood is safe from crime. |  |

### School Environment

The ‘School Environment’ variable is a validated subscale that was computed from the ABCD School Risk and Protective Factors Survey. The value of each item, listed below, was summed to obtain the final score. We obtained this score from the ABCD Sum Scores Culture & Environment measure (youth report).

| Item | Scoring |
| --- | --- |
| In my school, students have lots of chances to help decide things like class activities and rules. | 1=NO!, 2=no, 3=yes, 4=YES! |
| I get along with my teachers. |  |
| My teacher(s) notices when I am doing a good job and lets me know about it. |  |
| There are lots of chances for students in my school to get involved in sports, clubs, or other school activities outside of class. |  |
| I feel safe at my school. |  |
| The school lets my parents know when I have done something well. |  |

### Trauma Exposure

The ‘Trauma Exposure’ variable was computed by summing the total score per participant across the items below. This data was obtained from the ABCD Parent Diagnostic Interview for DSM-5 (KSADS) Traumatic Events measure (parent report).

| Item | Scoring |
| --- | --- |
| A car accident in which your child or another person in the car was hurt bad enough to require medical attention | 0=No, 1=Yes |
| Another significant accident for which your child needed specialized and intensive medical treatment |  |
| Witnessed or caught in a fire that caused significant property damage or personal injury |  |
| Witnessed or caught in a natural disaster that caused significant property damage or personal injury |  |
| Witnessed or present during an act of terrorism (e.g., Boston marathon bombing) |  |
| Witnessed death or mass destruction in a war zone |  |
| Witnessed someone shot or stabbed in the community |  |
| Shot, stabbed, or beaten brutally by a non-family member |  |
| Shot, stabbed, or beaten brutally by a grown up in the home |  |
| Beaten to the point of having bruises by a grown up in the home |  |
| A non-family member threatened to kill your child |  |
| A family member threatened to kill your child |  |
| Witness the grownups in the home push, shove or hit one another |  |
| A grown up in the home touched your child in his or her privates, had your child touch their privates, or did other sexual things to your child |  |
| An adult outside your family touched your child in his or her privates, had your child touch their privates or did other sexual things to your child |  |
| A peer forced your child to do something sexually |  |
| Learned about the sudden unexpected death of a loved one |  |

### Parent Support

The ‘Parent Support’ variable was computed as the mean of the items below from the ABCD Children’s Report of Parental Behavioral Inventory. We obtained this score from the ABCD Sum Scores Culture & Environment measure (youth report).

| Item | Scoring |
| --- | --- |
| First caregiver (caregiver participating in study/completing protocol). Makes me feel better after talking over my worries with him/her | 1 = Not like him/her; 2 = Somewhat like him/her; 3 = A lot like him/her |
| First caregiver (caregiver participating in study/completing protocol). Smiles at me very often. |  |
| First caregiver (caregiver participating in study/completing protocol). Is able to make me feel better when I am upset. |  |
| First caregiver (caregiver participating in study/completing protocol). Believes in showing his/her love for me. |  |
| First caregiver (caregiver participating in study/completing protocol). Is easy to talk to. |  |

### Caregiver Support

The ‘Caregiver Support’ variable was computed as the mean of the items below from the ABCD Children’s Report of Parental Behavioral Inventory. We obtained this score from the ABCD Sum Scores Culture & Environment measure (youth report).

| Item | Scoring |
| --- | --- |
| Second caregiver. Makes me feel better after talking over my worries with him/her. | 1 = Not like him/her; 2 = Somewhat like him/her; 3 = A lot like him/her |
| Second caregiver. Smiles at me very often. |  |
| Second caregiver. Is able to make me feel better when I am upset. |  |
| Second caregiver. Believes in showing his/her love for me. |  |
| Second caregiver. Is easy to talk to. |  |

### Family History of Mental Health Problems

The ‘Family History of Mental Health Problems’ variable was computed from data obtained from the ABCD Family History Assessments Parts 1 and 2 (parent report). This measure indexes whether family members including parents, siblings, aunts, uncles, cousins, and others have had mental health concerns, struggled with substance use, or been involved with criminal activities. We selected immediate family members (mother, father, and full siblings), then summed the number of endorsements for substance use, criminal activities, and mental health concerns per family member. We then summed the endorsements of each family member per participant.

| Item | Scoring |
| --- | --- |
| Has ANY blood relative of your child ever had any problems due to alcohol, such as: Marital separation or divorce; Laid off or fired from work; Arrests or DUIs; Alcohol harmed their health; In an alcohol treatment program; Suspended or expelled from school 2 or more times; Isolated self from family, caused arguments or were drunk a lot. | 0=No, 1=Yes |
| Has ANY blood relative of your child ever had any problems due to drugs, such as: Marital separation or divorce; Laid off or fired from work; Arrests or DUIs; Drugs harmed their health; In a drug treatment program; Suspended or expelled from school 2 or more times; Isolated self from family, caused arguments or were high a lot. |  |
| Has ANY blood relative of your child ever suffered from depression, that is, have they felt so low for a period of at least two weeks that they hardly ate or slept or couldn't work or do whatever they usually do? |  |
| Has ANY blood relative of your child ever had a period of time when others were concerned because they suddenly became more active day and night and seemed not to need any sleep and talked much more than usual for them? |  |
| Has ANY blood relative of your child ever had a period lasting six months when they saw visions or heard voices or thought people were spying on them or plotting against them? |  |
| Has ANY blood relative of your child been the kind of person who never holds a job for long, or gets into fights, or gets into trouble with the police from time to time, or had any trouble with the law as a child or an adult? |  |
| Has ANY blood relative of your child ever had any other problems with their nerves, or had a nervous breakdown? |  |
| Has ANY blood relative of your child ever attempted or committed suicide? |  |

### Parental Monitoring

The ‘Parental Monitoring’ variable was computed as the mean of the items below from the ABCD Parental Monitoring Survey. We obtained this score from the ABCD Sum Scores Culture & Environment measure (youth report).

| Item | Scoring |
| --- | --- |
| How often do your parents/guardians know where you are? | 1 = Never; 2 = Almost Never;<br>3 = Sometimes; 4 = Often; 5 =<br>Always or Almost Always |
| How often do your parents know who you are with when you are not at school and away from home? |  |
| If you are at home when your parents or guardians are not, how often do you know how to get in touch with them? |  |
| How often do you talk to your mom/dad or guardian about your plans for the coming day, such as your plans about what will happen at school or what you are going to do with friends? |  |
| In an average week, how many times do you and your parents/guardians, eat dinner together? |  |

### Family Conflict

The ‘Family Conflict’ variable is a validated subscale that was computed from the ABCD Youth Family Environment Scale-Family Conflict Subscale Modified from PhenX (FES). The value of each item, listed below, was summed to obtain the final score. We obtained this score from the ABCD Sum Scores Culture & Environment measure (parent report).

| Item | Scoring |
| --- | --- |
| We fight a lot in our family. | 1 = True; 0 = False |
| Family members rarely become openly angry. | 0 = True; 1 = False |
| Family members sometimes get so angry they throw things. | 1 = True; 0 = False |
| Family members hardly ever lose their tempers. | 0 = True; 1 = False |
| Family members often criticize each other. | 1 = True; 0 = False |
| Family members sometimes hit each other. | 1 = True; 0 = False |
| If there's a disagreement in our family, we try hard to smooth things over and keep the peace. | 0 = True; 1 = False |
| Family members often try to one-up or outdo each other. | 1 = True; 0 = False |
| In our family, we believe you don't ever get anywhere by raising your voice. | 0 = True; 1 = False |

